# A scalable method for parameter-free simulation and validation of mechanistic cellular signal transduction network models

**DOI:** 10.1101/107235

**Authors:** Jesper Romers, Sebastian Thieme, Ulrike Münzner, Marcus Krantz

**Affiliations:** Institute for Biology, Humboldt-Universität zu Berlin, Berlin, Germany

## Abstract

The metabolic modelling community has established the gold standard for bottom-up systems biology with reconstruction, validation and simulation of mechanistic genome-scale models. Similar methods have not been established for signal transduction networks, where the representation of internal states leads to scalability issues in both model formulation and execution. While rule- and agent-based methods allow efficient model definition and execution, respectively, model parametrisation introduces an additional layer of uncertainty due to the sparsity of reliably measured parameters. Here, we present a scalable method for parameter-free simulation of mechanistic signal transduction networks. It is based on rxncon and uses a bipartite Boolean logic with separate update rules for reactions and states. Using two generic update rules, we enable translation of any rxncon model into a unique Boolean model, which can be used for network validation and simulation - allowing the prediction of system level function directly from molecular mechanistic data. Through scalable model definition and simulation, and the independence of quantitative parameters, it opens up for simulation and validation of mechanistic genome-scale models of signal transduction networks.

## Introduction

Systems biology aims at the integrative analysis of large-scale biological systems up to whole cells. To realise this goal, we integrate knowledge into executable or computational models^1^. This process has been developed the furthest in the field of metabolic modelling, where the community routinely works with genome-scale models. These models are defined at the level of biochemical reactions, cover the entire metabolic network of even complex cells, and can be simulated to predict system level functionality^2,3^. The methodology is well established and supported by rich toolboxes for network reconstruction, validation and simulation^4^, and it constitutes the paradigm for bottom-up modelling. However, these tools cannot be used for signal transduction networks, due to the difference between mass and information transfer networks^5^.

Mechanistic modelling of signal transduction is challenging at several levels. First, at the level of model definition: Empirical data on site-specific modifications of, or bonds between, signalling components combine combinatorially into a large number of possible configurations, or microstates^6^. To solve this, the community developed simulation tools with adaptive resolution, such as the rule-based modelling languages BioNetGen and Kappa^7,8^. Second, simulation may be prohibitively expensive even with an efficient model definition, as is the case for classical rule-based modelling, in which the full network of microstates must be generated. This was solved by the development of the network-free simulation tool for rule based models, NFsim^9^. Third, quantitative dynamic models require rate laws that must be parametrised in terms of rate constants and initial molecule amounts. Reliable information on these quantities is sparse, precluding meaningful parametrisation of most mechanistic models. While the lack of quantitative knowledge is an experimental rather than a theoretical challenge (more dedicated biochemistry is needed), a simulation method that voids the need for parametrisation would be extremely helpful in evaluating mechanistic large-scale signalling models until this knowledge gap can be filled.

Parameter-free simulation of cellular networks is typically performed through constraint-based or Boolean methods^10^. Only the second is suitable for signal transduction, due to the lack of mass transfer through the network. In Boolean networks, nodes are either true or false and edges define how the node states at time t+1 depend on the node states at time t. Boolean networks have been used extensively in modelling, and different methods and toolboxes have been developed^11^. Most of these account for components but not their states, and hence omit the mechanisms of signal transduction. However, three mechanistic Boolean modelling methods have been developed: One is derived from SBGN-PD diagrams, one from rule-based models, and one is based on rxncon, the reaction-contingency language^12–14^. These methods support a detailed description of signalling events. However, the first is based on a microstate description, inheriting the scalability issues of these approaches^12^; the second requires a fully parametrised rule-based model, inheriting the problem of parametrisation^13^; and the third inherited shortcomings in expressiveness and precision from the first generation of the rxncon language^14^. Consequently, new methods are needed to support parameter-free, and hence scalable, simulation of signal transduction networks.

Here, we present a parameter-free simulation method that supports large-scale mechanistic models of signal transduction networks. This bipartite Boolean modelling (bBM) logic is based on the second generation rxncon language, which is tailored for formalising signal transduction models based on empirical data^15^: The reaction network is defined in terms of *elemental states*, i.e. modifications (or lack thereof) at specific residues and bonds (or lack thereof) at specific domains. The definition is bipartite: *Elemental reactions* are decontextualised reaction events that define how elemental states are *synthesised, degraded, produced* or *consumed*, and *contingencies* define how elemental reactions depend on (combinations of) elemental states that are not changed through the reaction. This structure is kept in the bBM: We developed two generic update rules, one for *elemental states* and one for *elemental reactions*, based on the behaviour of two minimal reaction motifs, and demonstrated that these motifs can be connected, LEGO-brick style, into large-scale models without any need for optimisation at the system level. Furthermore, we show that the resulting models meaningfully reproduce the behaviour of signalling processes, and that discrepancies between model behaviour and system levels expectations can be used to identify gaps in the network reconstruction and hence to improve the model. Taken together, we present a scalable method for parameter-free validation of mechanistic signal transduction network models, taking an important step to close the gap in capabilities between metabolic and signal transduction modelling, and introduce a method for scalable simulation of signal transduction networks that supports modelling at the genome scale.

## Results

### Generic update rules: basics

The central result presented in this work is a method to derive a parameter-free Boolean model from a rxncon network, in a completely mechanistic fashion.

A Boolean model exists of *nodes* or *targets* that carry a value - true or false - at a given point in time, and an update rule describing its value at the next time, formulated as a Boolean function of every node at the current time. The Boolean model we construct is *bipartite:* this means that there exist two classes of nodes, one class for reactions and the other for states. The interpretation we assign to the value of these nodes is as follows:

- If a *reaction node* is true, the cellular regulatory network is, at that point in time, in a configuration where it can accommodate that reaction. This is required, but not sufficient, for the reaction to take place: In the absence of its source state(s), a reaction will not “fire” even though its value is true, as the reaction nodes are purely a description of the regulatory layer of the biological cell.
- A *state node* is true if there are a sufficient number of molecules carrying that state present in the cell for it to be considered functionally relevant.

Both reaction and state nodes describe *system-level* properties, not molecular ones. A consequence of this is, for example, that two states that are mutually exclusive on a single molecule *(e.g*. a single residue being phosphorylated and unphosphorylated), can be simultaneously true in the Boolean system.

Reactions can act on states in four different ways: production, consumption, synthesis and degradation. The behaviour stems directly from the *skeleton rule* underlying the reaction^15^. This rule is similar to a rule-based model rule, such as can be defined in BNGL, but consists solely of a center, without context. We repeat the definitions in that work here, where RHS and LHS refer to the right-hand respectively left-hand side of the skeleton rule. *Production:* a state is produced by a reaction if it appears on the RHS, not on the LHS, but the component carrying the state does appear on the LHS. *Consumption:* a state is consumed by a reaction if it appears on the LHS, not on the RHS, but the component carrying the state does appear in the RHS. *Synthesis:* a state is synthesised by a reaction if it appears on the RHS, and the component carrying the state does not appear on the LHS. *Degradation:* a state is degraded by a reaction if the component carrying the state appears on the LHS, no state mutually exclusive with it appears on the LHS, and the component carrying the state does not appear on the RHS.

In what follows, we study the smallest irreducible motifs containing two states (for modification reactions; Fig 1) or three states (for interactions; Fig S1), and the reactions acting upon them. We define, given an initial configuration for the states and the absence or presence of each of the reactions, the desired steady-state behaviour. The crux of the matter then becomes to find update rules for the states that reproduce this steady state. Our desired behaviour originates in experiment, and is based on the observation that there exists a natural hierarchy between the reaction types introduced in the previous paragraph: *synthesis* is stronger than *degradation* which is stronger than *production* which is stronger than *consumption*.^1^

**Figure 1:**
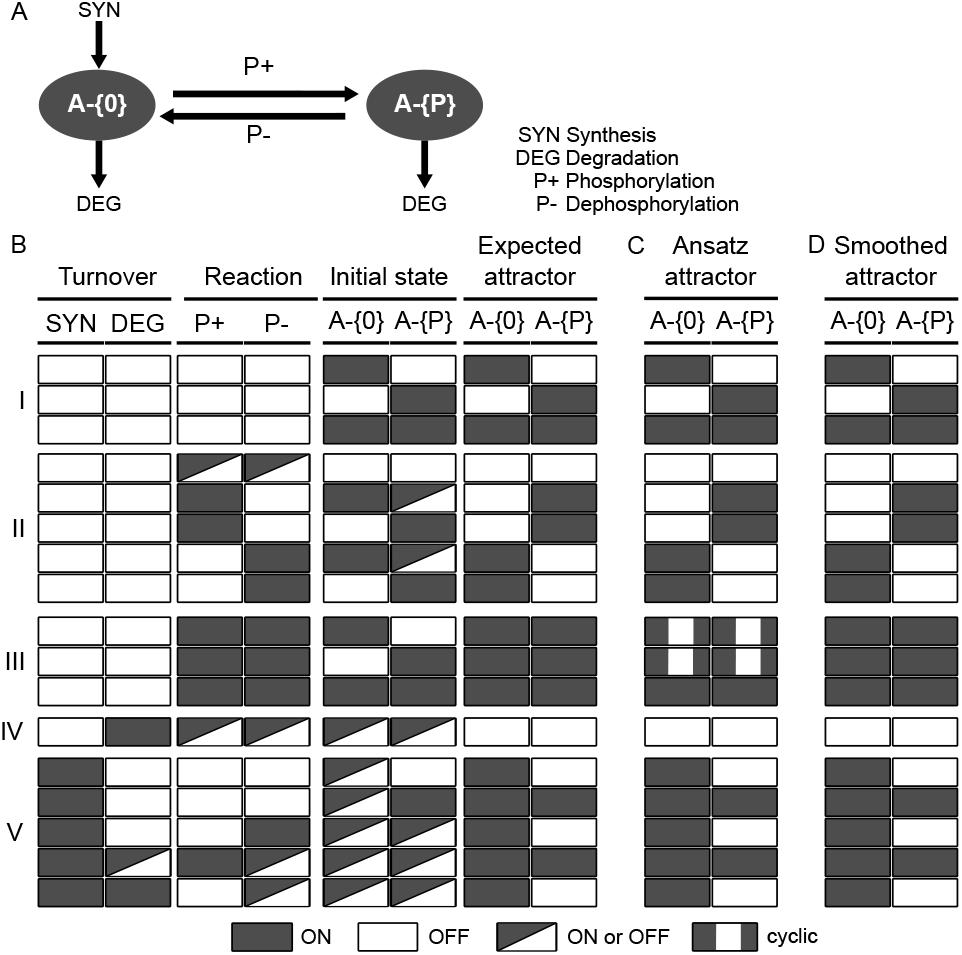
Behaviour of a minimal modification motif. **(A)** The motif includes two states of A, unphosphorylated (A-{0}) and phosphorylated (A-{P}). The motif contains up to four different reactions: Component A can be synthesised (in its neutral state A-{0}), degraded (in either state), phosphorylated (consumes A-{0}, produces A-{P}) or dephosphorylated (consumes A-{P}, produces A-{0}). **(B)** The expected steady state as a function of initial state and active reactions. (I) In the absence of any reactions, the equilibrium state will be identical to the initial state. (II-III) In absence of synthesis or degradation, but in the presence of component A (A-{0} or A-{P} true), the equilibrium depends on the (de)phosphorylation reactions. With only one of these reactions, only the fully (de)phosphorylated form is present. However, with both reactions present we expect both elemental states to be present at steady state. (IV) With degradation but not synthesis active, the protein will be depleted and both states will be false. (V) With active synthesis, the neutral state will always be present. The phosphorylated state will only be present if there is also a phosphorylation reaction or if the state is initially present and both the degradation and dephosphorylation reactions are off. **(C)** The simulation outcome with the update rules in the original ansatz. The results correspond to the expected behaviour for 62 out of 64 configurations. The exception are the cyclic attractors in block (III), where both phosphorylation and dephosphorylation are active and only one of the elemental states are initiated as true. **(D)** The simulation outcome with the smoothed update rules. The results are identical to the expected attractor.

### Defining the notation: reactions, states and components

A rxncon system will contain *N_R_* reactions denoted by *R_i_, N_s_* elemental states denoted by *S_i_*, and *N_c_* components denoted by *C_i_*. For the components that appear without any internal states, such as e.g. those that solely appear as kinases, the component is its own state (which does not appear in the original rxncon system). For those components that do carry internal states, the component can be expanded as a Boolean expression of elemental states grouped by the site (domain or residue) on which they live:

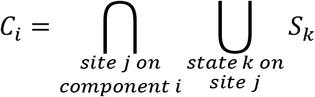

The origin of this expression is in the mutual exclusivity of states that live on the same residue or domain: for each of these sites, at least one of the states living on the site needs to be present for the component itself to be present. A consequence of this is that for components that carry internal states, the dynamic behaviour of the component is fully determined by the states which the component carries.

Furthermore, we define functions mapping states and reactions to Boolean expressions of states. First, the functions *K(R_i_)* list the components which are reacting in reaction *R_i_*:

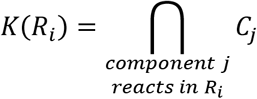

Whereas the functions *K(S_i_)* list the components carrying the state *S_i_* (bond states are carried by two components, whereas modifications are carried by one):

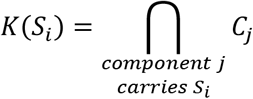

In the update rule for the states, we will require the following combinations. First, a reaction together with its source states:

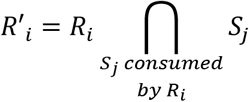

The neutral state (unmodified, unbound) counterpart for a particular state *S_i_* is denoted by *N(S_i_)*.

These notations enable us to write down the synthesis term Σ, which describes whether a component is either directly or indirectly being synthesized:

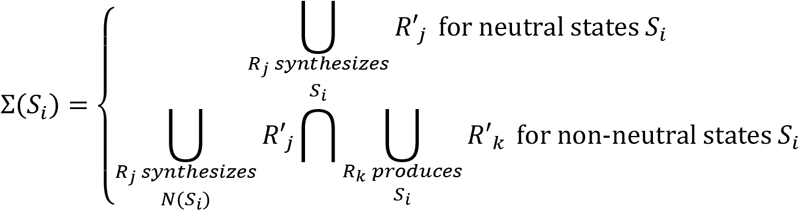

All systems we considered contain only states “one step” removed from the neutral states, so the expression for non-neutral states describes an active path coming from the synthesized state to the state under consideration. For states that are multiple steps removed from the synthesized neutral state, this expression has to be appropriately amended.

Finally, we denote the Boolean expression representing the contingency for reaction *R_i_* by *L(R_i_)*:

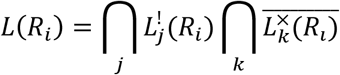

where the *L^!^_j_(R_i_)* and *L^x^_k_(R_i_)* enumerate the required respectively inhibitory contingencies for reaction *R_i_*, which are themselves possibly nested Boolean expressions.

### The expected behaviour of a small reaction circuit and update rule ansatz

The reaction update rules are quite straightforward. We require the strict contingencies^2^ for the reaction to be satisfied and the presence of the components on which the reaction acts.

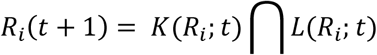

The state update rules are more complex and require explicitly the hierarchy between types of reactions that was alluded to above. First of all, if a state is synthesised by any reaction, it will be true. If synthesis is false, the necessary, but not sufficient requirement for the state to be true is that degradation is also false, and that the component(s) carrying the state are present. Now, there are two options for the state to be true: either the state is being produced by some reaction, in which case it is immaterial what the previous value of the state was, or the state was already true and it is not being consumed by any reaction (Fig 1, Fig S1).

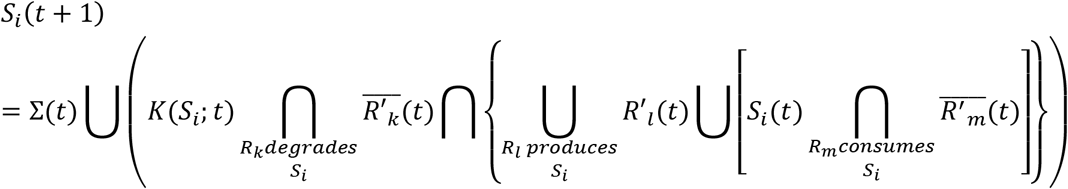

As can be seen in this formula, the translation of “the state is being produced” *et cetera* contains the primed reactions, meaning the reaction producing that state and the source state(s) of that reaction. This is due to the semantics of the reaction nodes, which only tell us about the regulatory state of the network.

### Testing the generic update rules

To test if the update logic described above captures the expected behaviour, we implemented the model generation process (see methods) and used it to generate the 64 models corresponding to the minimal modification circuitry above (Fig 1A). The models were simulated using BoolNet^16^, as described in the methods. The attractor states are visualised in Figure 1C. The model behaviours correspond to our expectations with one notable exception. In the absence of synthesis and degradation, but in the presence of both phosphorylation and dephosphorylation, the model displays an oscillatory behaviour when only one of the two states is initiated. Closer inspection reveals that this is due to periodic source state depletion. The phosphorylation and dephosphorylation reactions are constitutive (no contingencies, no loss of components), and the oscillations due to the state updates, which are completely encoded in the state update rule. Indeed, as soon as a reaction executes, dependent on its source state, it depletes the source state pool. Hence, the reactions alternate in firing, triggering out-of-phase oscillations in the truth value of the states. Consistently, these oscillations disappear when both states are initiated or when the source state is repleted through synthesis. We observe the same phenomenon in the interaction motif, when the reaction cycle is active and at least one component is initiated in and remains in a single form (Fig S1C). We consider these spurious oscillations undesirable in our systems-level description, but note that they would be appropriate for models of single molecules. Nevertheless, the outcome is highly encouraging, as 62 out of 64 models matched the expected behaviour.

### Source state smoothing eliminates the spurious oscillations

To eliminate the oscillations that plagued our initial ansatz, we adapted the state target updates by widening the window in which we checked for source state availability: A reaction needs the source state to be present, or to be produced. To achieve this, in formula (8) we substitute for the reactions producing the state, *R’_l_* the following:

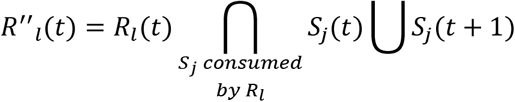

Where the Sj(t+1) is the full expression (8), without this substitution.

We consider this adaptation quite natural: there are large numbers of molecules undergoing the same set of reactions. Therefore, it is highly unlikely that all of these reactions are temporally completely in phase, justifying a smoothing over molecules. In addition, the time scale in a Boolean model is basically set by the slowest of the reactions, since all rate constants are absent. For molecule pools acted upon by reactions that are faster than the slowest in the system, it is likely that they will pass through mutually exclusive states within the window of a Boolean time step, justifying a “time smoothing”. Both effects are captured by the reworked update rules.

The smoothing assumption only breaks down in the context of few molecules and low reaction rates, and there are cases in which smoothing is inappropriate. In other work^17^, we have introduced a hybrid model containing both the molecular reactions and states described here, and additionally macroscopic reactions and states that are governed by the non-smoothed update rules. However, for most states in a signal transduction network, the smoothed update rule is more appropriate. Having established the smoothing logic, we implemented it into the rxncon compiler tool (see methods) and recreated the 64 models with smoothing. We repeated the simulation and compared the results to the original simulation (Fig 1D, Fig S1D). The oscillatory behaviour disappeared, but no other simulation results changed. Hence, the simulation results exactly match the behaviour we expect from a model of these reaction motifs.

### The update rules can be used as LEGO bricks to assemble a systems level model

Next, we applied the bBM logic to simulate a linear pathway. We chose a simplified model of the HOG MAP kinase pathway from *Saccharomyces cerevisiae* (Fig 2; taken from^14^). We created a rxncon 2.0 model of this pathway (Table S1), and used this to generate the bBM using the generic update rules with smoothing. Already this small model has 28 reaction and state targets, and hence 2^28^ (~10^8^) possible initial states. We deemed this too many for an exhaustive search, and decided to use a generic start state for all simulations: All neutral elemental states (that is, unbound binding domains and neutral modifications) are true, all generic component states (for components with no elemental states) are true, and all other nodes are false. From this highly artificial initial state, we let the model find its own natural “off-state” by executing it until an attractor is reached (Fig 2B). At this point, we change the input state and repeat to see the response of the pathway to the input, and repeat this process until the model returns to a state we have already seen. As can be seen from Figure 2B, the HOG pathway responds appropriately to turgor: It turns off the kinase cascade. For comparison, we repeated the simulation with the non-smoothed logic (Fig S2A), where we see the signal passing through the network despite spurious oscillations. However, the system does not converge to point attractors, leading to more complex analysis and interpretation. There are three striking blocks in the heat-map (Fig 2B): First, the initial neutral states never turn off. Second, there is a block of reactions that turns on directly, and stays on throughout the simulation. Third, there is a block that turns on and off in response to the signal. The third block contains the reactions and states that actually transmit the information. The second block contains constitutive reactions, which are either unregulated (e.g. dephosphorylation reactions), or regulated at the level of source state availability (e.g. phosphotransfer from Sln1 to Ypd1). The first block contains all the neutral states. These remain true because the reactions that produce them are considered unregulated, which may be due to experimental bias as discussed below. Hence, the logic of the generic update rules is sufficient to convert the molecular level knowledge in a rxncon network into a functional bBM that accurately predicts system level function. It is highly non-trivial that generic update rules, which were defined for isolated reactions, suffice to define a complete model that functions at the systems level, with no further tweaking or parametrisation. Taken together, the generic update rules map any given rxncon network on a unique Boolean model that predicts systems level function.

**Figure 2:**
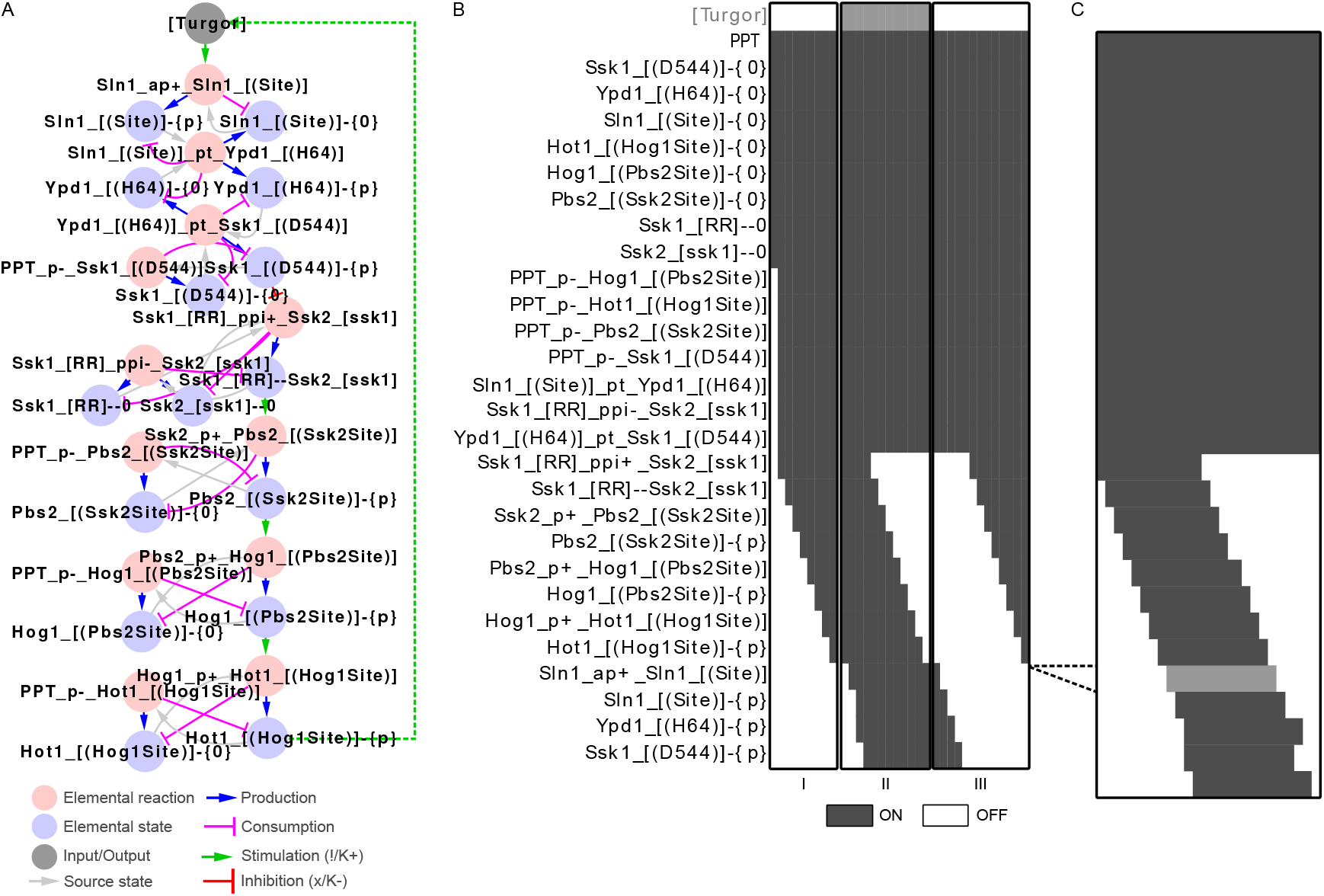
From reactions to a functional pathway. We used the smoothed update rules to generate and simulate a model of the Sln1 branch of the high osmolarity glycerol (HOG) pathway. (A) The pathway visualised as a rxncon regulatory graph. In the absence of turgor, Sln1 stays unphosphorylated. As turgor increases, the auto-phosphorylation of Sln1 initiates a phosphotransfer cascade converging at Ssk1. The phosphorylated form of Ssk1 turns off the downstream MAP kinase pathway leading to dephosphorylation of the downstream transcription factor Hot1. The dashed line indicates a feedback loop that is included in the cyclic version only. (B) Simulation of a linear version of the model using source state smoothing of the update rules. (I) We use our default assumptions on the initial state and simulate the model until we reach an attractor (first OFF trajectory). (II) We activate the system by turning [Turgor] ON and simulate again (ON trajectory) until we reach an attractor state. (III) From there, we set [Turgor] OFF again and simulate the model until we reach an attractor. We observe that the model responds as expected to the input. (C) We extended the HOG model with a feedback loop, where activation of the pathway leads to increased turgor (via Hot1-{P}). This is simplification of an adaptive response through increased glycerol production and retention, which increases turgor. We simulate this model from the initial OFF attractor (see panel B), and note that the system oscillates as expected: The trajectory is cyclic, where the last time step is followed by the first, and turgor is now a model variable that turns on and off during the cycle (grey row in the heatmap).

### The bipartite Boolean logic correctly reproduces real oscillations

The HOG pathway is a homeostatic pathway that maintains proper turgor pressure. The pathway output eventually leads to signal cessation through a physiological feedback loop^18^. To simulate this, we linked the most downstream component to the input that turns the pathway off (dashed line in Fig 2A). We repeated the model creation and simulation, using the initial steady state of the linear model as starting condition. As shown in Figure 2C, the model now shows a periodic activation/deactivation behaviour, similar to that when the input is changed manually. Hence, the bBM logic is fully capable of predicting biologically relevant oscillations. As comparison, we also simulated the cyclic HOG model without source state smoothing (Fig S2B). Here, the pathway signal is completely washed out by the spurious oscillations and breaks down to a two-state cyclic attractor. The source state smoothing facilitates bBM analysis and clearly improves the interpretability of the simulation results. Taken together, the bBM logic generates Boolean models that can predict systems level function for both linear and cyclic systems.

### The bipartite Boolean logic scales to large-scale systems

Finally, we applied the method on the pheromone response pathway of baker’s yeast. We chose this pathway to benchmark the bBM method due to the existence of an excellently annotated, comprehensive and mechanistically detailed rule based model (RBM)^19^. The original RBM contains 229 rules with 200 parameters (166 unknown) that define how 18 components can assume over 200.000 distinct states (http://yeastpheromonemodel.org/wiki/Extracting_the_model). While this is one of the most carefully built and curated RBMs, it remains difficult to meaningfully simulate it as such^20^. Hence, it constitutes an excellent benchmark target for the bBM method.

We simulated the pheromone bBM using a standardised simulation workflow (see methods). The rxncon translation of the RBM, which is described elsewhere^15^, is defined by 95 elemental reactions and 231 (non-zero) contingencies. We generated the bBM using the smoothed update rules, which produced a bipartite Boolean model with 130 reaction targets (due to separation of bidirectional reactions and duplication of degradation reaction targets) and 118 state targets. With 248 state variables, the model is too large to use an exhaustive search of initial states (statespace = 2^248^; ca 10^74^ distinct configurations), so we rely on the default initiation state (all neutral state targets are true, all generic component targets true, all other targets are false). From this initiation state, we first let the model find its natural “off-state”, as explained for the HOG pathway above, before we iteratively switched the input to true and false. We found that the pathway was constitutively active and unresponsive to pheromone. First, we examined if this was due to the interpretation of quantitative effects that are lost in the Boolean model. However, neither ignoring all nor including all K+/K− contingencies solves the problem. Furthermore, the original RBM was never simulated and proven functional. Hence, we proceeded with the minimal model (ignoring quantitative contingencies) and looked deeper into the pathway behaviour, finding that it activates in the absence of signal due to constitutive release of Ste4, which represents the beta/gamma subunit of the trimeric G-protein at the top of the cascade. To address the problem, we change two (out of 87) quantitative contingencies into qualitative contingencies, indicating that a much simpler model would suffice to capture the key features of the pathway. In addition, we needed to limit turnover of Ste4 bound Gpa1 (to prevent signal-independent release of unbound Ste4, the activator of the pathway) and to remove Fus3 dependent degradation of Ste12 which made the pathway “single-shot”. The final model with changes can be found in Table S2, and the simulation trajectories are shown in Figure 3. This updated version of the model responds to pheromone exposure and withdrawal as expected.

**Figure 3:**
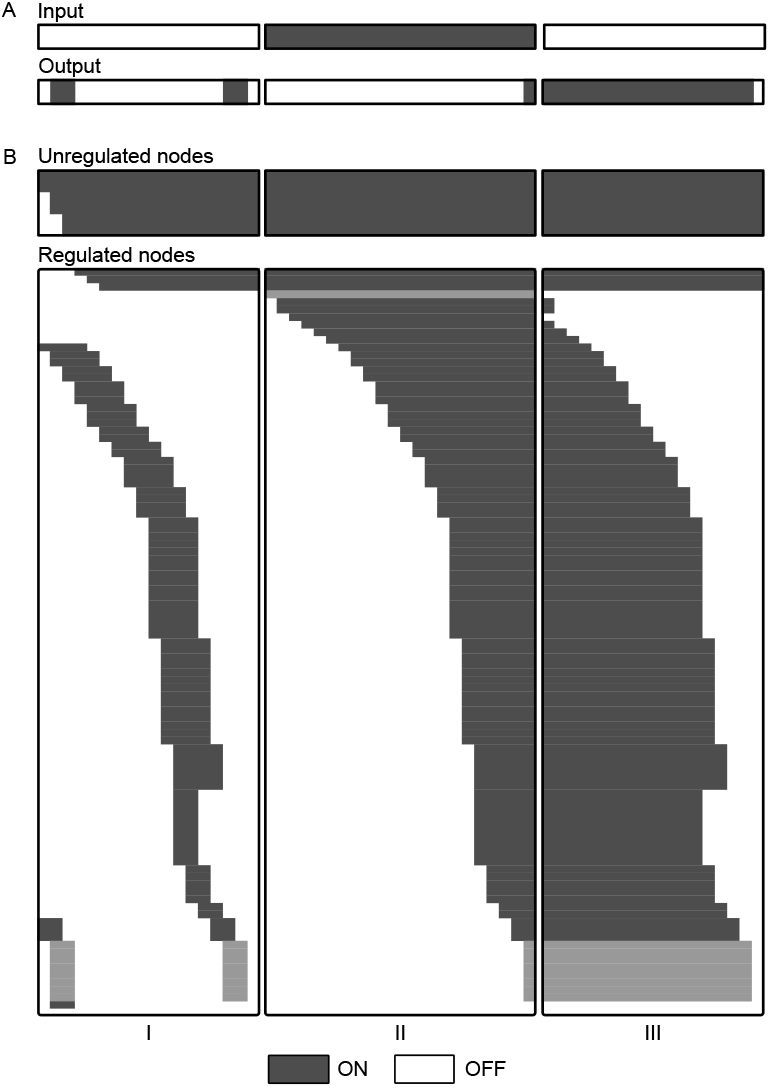
The updated pheromone model responds as expected to pheromone. We generated a bBM form the updated pheromone model and (I) simulated it in the absence of pheromone (unbound pheromone set to false) until the first steady state, where (II) free pheromone was set to true, representing pheromone stimulation, and the simulation repeated until next steady state was reached, before (III) pheromone was removed (by setting both free and bound pheromone to false) and the model simulated to the next steady state. The pathway turns on and off as expected, and finds a natural off state in the first simulation despite two activation pulses that go through the pathway. These are due to proteins that activate the pathway in their neutral states: The upstream Ste4, as discussed in the text, and the Ste12 transcription factor - which, according to the model, is active when unbound to Dig1 and Dig2. However, both pulses are transient and the steady state is robust against these transient dynamics.

## Discussion

Here, we present a qualitative simulation method for large-scale, mechanistically detailed signal transduction network models. The formalism is based on Boolean logic and can be simulated and studied by a standard package such as BoolNet. However, we present a fundamentally new concept to formulate Boolean models. First, we create a bipartite model at the level of elemental reactions and contingencies, capturing the key elements in signal transduction at an appropriate resolution for mechanistic modelling of these processes. Second, based on detailed analysis of two minimal reaction motifs, and on a small set of standard assumptions, we define two generic update rules: one for reaction and one for state targets. These generic update rules map a bipartite rxncon network on a unique bBM with defined truth tables. The elemental reactions define the update rules for state targets, and the contingencies define the update rules for reaction targets. We show that these building blocks can be assembled like LEGO-bricks into a bipartite Boolean model that predicts system level function from molecular mechanisms, without optimisation at the system level.

The unique mapping from rxncon to an executable bBM that predicts system behaviour is highly non-trivial. Normally, it is relatively easy to build a Boolean model structure, but highly non-trivial to define truth tables that enable the model to reproduce the behaviour of the system^21^. Here, we find that the regulatory structure encoded in the rxncon network already uniquely defines a Boolean model with set truth tables, and that this Boolean model meaningfully predicts system level behaviour. Thus, the bBM logic we present here bridges the microscopic (biochemical reactions) and macroscopic (input-output) levels of cellular signal transduction, fulfilling the requirements for the cellular “mechanics” proposed by Hlavacek and Faeder - at least qualitatively^6^.

This has far-reaching implications: First, it provides an efficient validation tool in the model building process. This allows the model construction to be separated in to two phases: A qualitative and a quantitative phase. Boolean models are computationally inexpensive, and the automatic model generation supports iterative model creation, analysis and improvement. In addition, we are better equipped with knowledge at the qualitative level, suggesting that this level should be optimised first. As rxncon supports compilation into both RBMs and bBMs (as well as several graphical formats), it can be used to facilitate this process^22^: The structural model can be created and validated using graphical tools and bBM simulation, and later the improved network can be used to create a rule-based model. Hence, the more expensive parameterisation cycles can be performed after the qualitative model has passed the validation process. Second, the bBM method can be used for validation of large-scale signal transduction networks. Previously, large-scale reconstruction of signal transduction has been limited to graphical maps that cannot be executed^23–25^. The method we present here changes this: We can now validate – through simulation – large-scale reconstructions of signal transduction networks.

The bBM formalism we presented here is fundamentally different from its previous incarnation^14^. First, it has been designed to capture system level behaviour, not the states of individual molecule instances in the network, meaning that states that would be mutually exclusive on a single molecule can be true at the same time. As we show in figure S2, this is critical for meaningful system level predictions. Second, we used a constructive approach, defining all possible behaviours at the level of two families of minimal reaction motifs, to design two generic update rules. These update rules can be used to map any rxncon system, even extended by new reaction types, through the interpretation of the flexible skeleton rule definition as synthesis, degradation, production, or consumption of different states. Third, the method we present here inherits the expressiveness and flexibility of rxncon 2.0, including the explicit representation of neutral states^15^. Fourth, the reimplementation has improved the model generation, enabled the use of different export options, and improved the model creation and analysis workflow. While the previous incarnation worked well in many instances^14,26,27^, these models had issues with certain reaction types (most notably degradation) and spurious oscillations. That the latter appeared so rarely was due to the implicit dominance of modified states: Neutral states were not explicitly represented, making modification or binding reactions dominant over reactions that returned components to their neutral state. Here, we eliminate this artificial hierarchy, which we consider undesirable, and explicitly include the neutral states. However, most of these states are constantly true in our simulations. This may have two reasons: First, it could reflect biology: There would be a constant pool of unmodified components as long as there is a constant turnover (and hence synthesis, which per definition occurs in the neutral states). Second, it could reflect an experimental bias, as we know much more about the modifying reactions than about the reactions that reverse the modification (e.g. phosphorylation vs dephosphorylation,^28^). If so, the formalism we present here helps us make this information bias explicit, and will allow us to integrate the regulation on these reactions as the knowledge becomes available.

Finally, the method enables a scale-shift in signal transduction modelling. Hitherto, executable signalling models have been mechanistically detailed or large-scale, but not both. Most mechanistic large-scale reconstructions are technically microstate models that could be simulated after parametrisation. However, they are actually divided into several unconnected modules and could hence not be simulated at the system level^23–25^. Rule-based modelling languages have been used to build relatively large models^20,29^, but even these models are limited to few (18 in these cases) components and parametrisation is already an outstanding challenge. In contrast, comprehensive signalling models will need to account for hundreds or thousands of signalling components, carrying many thousands of distinct elemental states. Here, we present a method that can deal with mechanistic signalling networks at this scope, and we have successfully used this method to build and analyse a comprehensive mechanistic model of the yeast cell division cycle, which accounts for 229 proteins, 790 elemental reactions and 1238 elemental states down to residue resolution when applicable^17^. The method is qualitative, but this may be an advantage given the sparsity of reliable quantitative information on rate constants. In addition, even metabolic modelling – clearly the state-of-the-art in genome-scale modelling – is limited to qualitative or semi-quantitative simulation methods at the genome scale.

Taken together, we present a parameter-free model creation and simulation method that enables simulation of genome-scale models of signal transduction. Together with the rxncon 2.0 language, it offers the possibility to build, validate and simulate genome-scale models of signal transduction networks – that can be turned into rule based models as soon as the quantitative knowledge makes it meaningful.

## Acknowledgements

This work was supported by the German Federal Ministry of Education and Research via e:Bio Cellemental (FKZ0316193, to MK).

## Appendix: Methods

### Model creation and analysis

The creation of the rxncon models are described elsewhere. The High Osmolarity Glycerol (HOG) model was taken from^14^ and adapted to rxncon 2.0. The pheromone (PHER) model was translated from the yeastpheromonemodel.org wiki as described in^15^. The bipartite Boolean model files were created with the rxncon compiler software, by calling the “rxncon2boolnet.py” script with default setting. The rxncon software is open source, distributed under the lGPL licence, and can either be downloaded from https://github.com/rxncon/rxncon or installed from the python package index with “pip install rxncon”. The rxncon model files are available as Table S1 (HOG model), Table S2 (final PHER model) or through download from https://github.com/rxncon/models/ (initial PHER model; YeastPheromoneModel.xls).

The rxncon2boolnet.py script generates three files. First, the bipartite Boolean model file (<ModelName>.boolnet) contains the update rules using states and reaction IDs. Second, the symbol mapping file (<ModelName>_symbols.csv) defines which IDs correspond to which states and reactions in the rxncon file. Third, the initial vector (<ModelName>_initial_vals.csv’) sets the initial state of the Boolean simulation.

Model simulation was done with the R CRAN package BoolNet^16^. To facilitate simulation, we prepared an R script that can be downloaded from https://github.com/rxncon/tools (BoolNetSim.R). To use this script through R studio:

- Save the network files and the R script into a single directory.
- Start RStudio.
- Open a new project and create it in the directory where you saved your files.
- Make sure your model files are located in the project folder.
- Open the R script. Set the filePrefix in the R script to <model>.
- Execute the entire script by selecting all text (ctrl+a) and pressing ctrl+enter.

The script generates five files: (i) <ModelName>.pdf, which graphically displays the simulation trajectory from initial state to the attractor, (ii) <ModelName>_trajectory_first.csv, with the trajectory as values (0/1) in tabular format, (iii) <ModelName>_2.pdf, which graphically displays the simulation trajectory from the attractor (useful to distinguish a point attractor (two columns) from a cyclic attractor (>2 columns), (iv) <ModelName>_trajectory_second.csv, with the second trajectory in tabular format, and (v) <ModelName>_new_attractor.csv, with the new attractor as an initial values file.

Where “<ModelName>” is the file name (without extension) of your rxncon model.

Within a Boolean model, we expect the output to be responsive to the input. The sign of the dependence does not matter, and we can start with the input either on or off. We simulate the model until it reaches an attractor. If it is a point attractor, we use it as starting point for the next simulation but turn the Input signal into a truth value, activating the output target and simulate again until we reach another attractor. We iteratively change inputs and simulate to an attractor state until we reach an attractor we have already seen.

---

**Figure S1:**
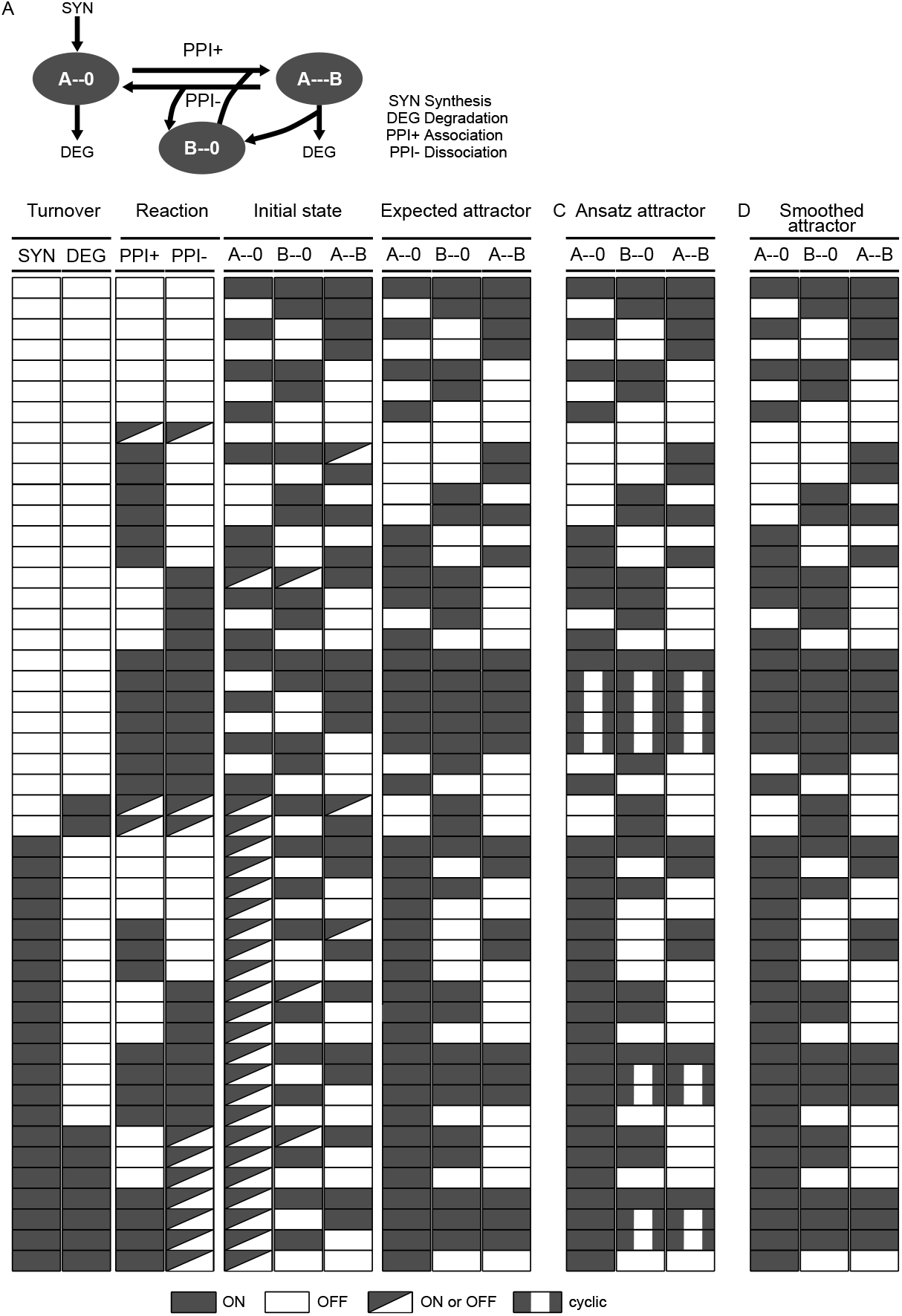
Behaviour of a minimal interaction motif. **(A)** The motif includes the unbound state of A (A--0), the unbound state of B (B--0) and the bound state (A--B). As in figure 1, the motif contains up to four different reactions: Component A can be synthesised (in its neutral state A--0), degraded (in either state), A and B can bind (ppi+; consumes A-- 0 and B--0, produces A--B) or dissociate (ppi-consumes A--B, produces A-- 0 and B--0). Not that degradation of A in the A-- B dimer releases B--0, hence this reaction is a degradation reaction of A and conditional production reaction for B--0. **(B)** The expected steady state as a function of initial state and active reactions. As in figure 1, the initial state is preserved when no reaction is active. However, the steady state in the presence of reactions is more complex, as both unbound states are necessary for the reaction to fire - which affects both the generation of the A--B state and the depletion of the unbound states. With degradation and without synthesis, A--0 and A--B is removed, releasing B--0 in the latter case. Finally, synthesis of A only leads to A--B in the presence of the forward reaction, in which case B--0 is depleted unless A--B is turned over by degradation or dissociation. **(C)** The simulation outcome with the update rules in the original ansatz. The results correspond to the expected behaviour, except when a component is present in only one state. This happens in the analogous case to the spurious oscillations in the modification motif in figure 1, but also when A is synthesised if B is only present in one form. **(D)** The simulation outcome with the smoothed update rules. The results are identical to the expected attractor.

**Figure S2:**
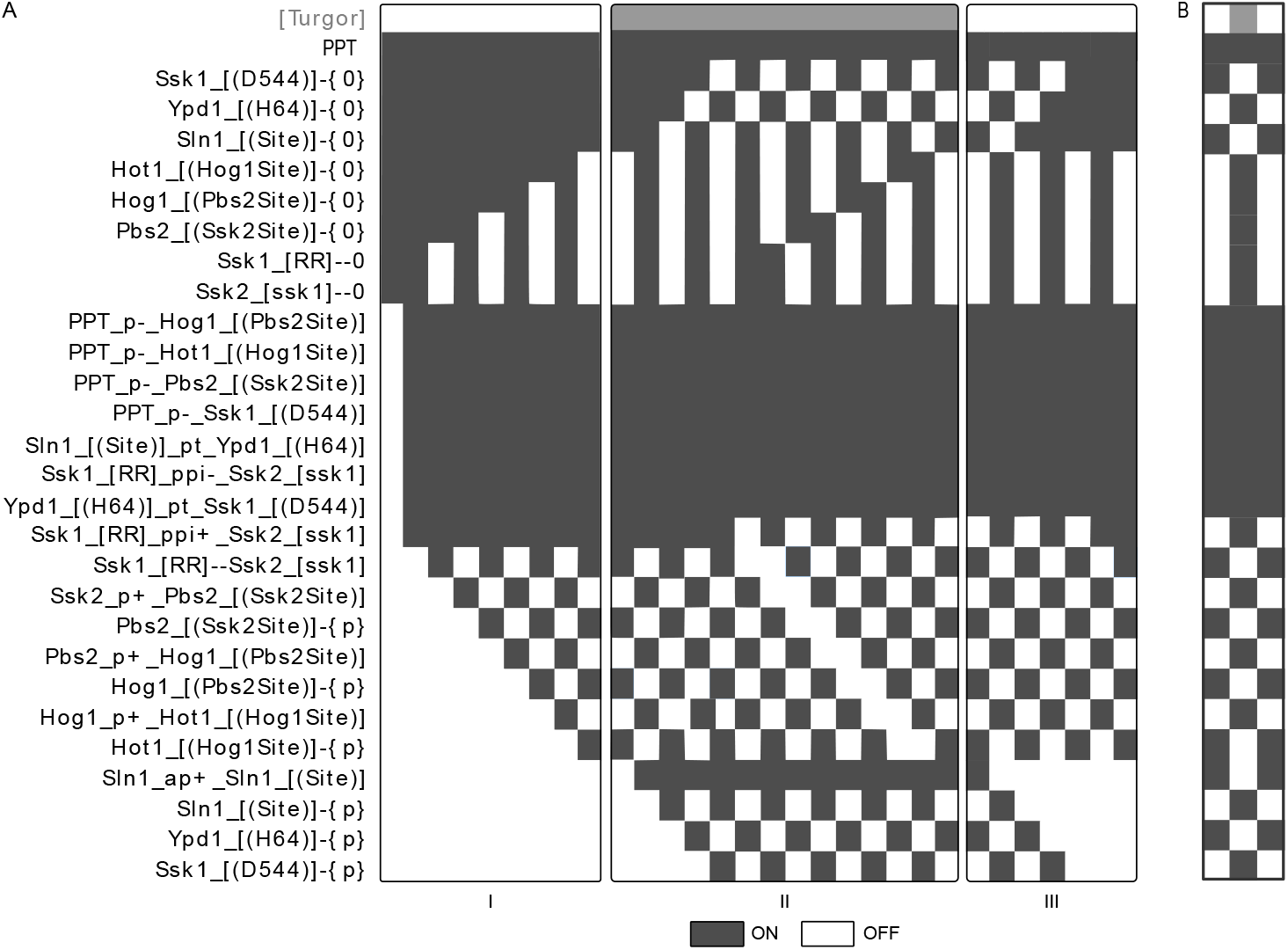
Smoothing is required to make pathway simulation interpretable. We repeated the simulation of the HOG pathway with the un-smoothed update rules from the initial ansatz. (A) Simulation of the linear model without smoothing. The signal goes through the pathway, but analysis is complicated by the spurious oscillations as the system no longer converges on point attractors. Each of the three simulation trajectories (I-III) ends with a cyclic attractor of length two. (B) Simulation of the cyclic HOG model without smoothing. Here, the entire oscillation cycle breaks down into a period two oscillator involving all the states (and their complements) and reactions that transmit the information.

**Table S1: The Hog pathway model in the rxncon format (acyclic)**.

**Table S2: The pheromone pathway model in the rxncon format**.

## Footnotes

1 Due to the crude concept of Boolean time, we consider a quasi-steady state at each update step. It is then easy to see that a protein that is both synthesised and degraded must be present in the system. In the case where the synthesis reaction is too weak to maintain functional level of the protein, it would be considered off. Similar, production, from the perspective of a specific state, will be dominant over consumption: In the presence of a phosphorylation cycle with both kinases and phosphatases active, both forms will be present. Finally, degradation is dominant over production, as depletion of a protein will deplete the phosphorylated form regardless of kinase activity (except when protected from degradation, as covered in the contingencies below).

2 Quantitative contingencies are special: in a Boolean system the reaction rate cannot be increased or decreased. In our analyses we choose to ignore them.

